# Changes in adenosine and neurotrophic signaling in the SOD1^G93A^ mouse model of amyotrophic lateral sclerosis: modulation by chronic caffeine intake

**DOI:** 10.1101/2022.07.14.500069

**Authors:** Nádia Rei, Cláudia A. Valente, Sandra H. Vaz, Miguel Farinha-Ferreira, Joaquim A. Ribeiro, Ana M. Sebastião

**Author notes:** **Correspondence:** Ana M. Sebastião, Instituto de Farmacologia e Neurociências, Faculdade de Medicina, Universidade de Lisboa, 1649-028 Lisboa, Portugal.

## Abstract

Amyotrophic lateral sclerosis (ALS) is characterized by the progressive degeneration of corticospinal tract motor neurons. Previous studies showed that adenosine-mediated neuromodulation is disturbed in ALS and that vascular endothelial growth factor (VEGF) has a neuroprotective function in ALS mouse models. We evaluated how adenosine (A_1_R and A_2A_R) and VEGF (VEGFA, VEGFB, VEGFR-1 and VEGFR-2) system markers are altered in the cortex and spinal cord of pre-symptomatic and symptomatic SOD1^G93A^ mice. We then assessed if/how chronic treatment of SOD1G93A mice with a widely consumed adenosine receptor antagonist, caffeine, modulates VEGF system and/or the levels of Brain-derived Neurotrophic Factor (BDNF), known to be under control of A_2A_R. We found out decreases in A_1_R and increases in A_2A_R levels even before disease onset. Concerning the VEGF system, we detected increases of VEGFB and VEGFR-2 levels in the spinal cord at pre-symptomatic stage, which reverses at the symptomatic stage, and decreases of VEGFA levels in the cortex, in very late disease states. Chronic treatment with caffeine rescued cortical A_1_R levels in SOD1^G93A^ mice, bringing them to control levels, while rendering VEGF signaling nearly unaffected. In contrast, BDNF levels were significantly affected in SOD1^G93A^ mice treated with caffeine, being decreased in the cortex and increased in spinal the cord. Altogether, these findings suggest an early dysfunction of the adenosinergic system in ALS and highlights the possibility that the negative influence of caffeine previously reported in ALS animal models results from interference with BDNF rather than with the VEGF signaling molecules.

## Introduction

Amyotrophic lateral sclerosis (ALS) is one of the most devastating neurodegenerative disorders, being the most common form of Motor Neuron Disease (MND). Both the upper (projecting from the cortex to the brainstem and the spinal cord) and lower motor neurons (projecting from the brainstem or spinal cord to the muscle) degenerate and ultimately die.

While ALS is a complex multifactorial disease, in which the pathophysiological mechanisms underlying motor neuron degeneration remain incompletely known, it has been established that adenosine-mediated neuromodulation is altered in this disease.^1^ Adenosine is a ubiquitous endogenous neuroprotective agent, involved in several biochemical processes, and plays a crucial role as a neuromodulator of synaptic transmission and plasticity, in both central and peripheral nervous systems. It functions through the actions of four G-protein coupled receptors – A_1_R, A_2A_R, A_2B_R and A_3_R, of which the A_1_R and A_2A_R are the most relevant, given their greater affinity for adenosine. While A_1_R are G_i/o_ coupled, leading to predominantly inhibitory effects, A_2A_R are G_s_ coupled, having an excitatory effect when activated. Congruently with the widespread involvement of the adenosinergic system in other neurodegenerative disorders, several studies have shown increased adenosine levels in the cerebrospinal fluid of ALS patients^2^, as well as increased A_2A_R expression in the lymphocytes^3^ and post-mortem spinal cord samples of ALS patients^4^.

Like adenosine, Vascular Endothelial Growth Factor (VEGF) has been associated to ALS. This system is comprised of three receptors (VEGFR-1/2/3), all of which are tyrosine kinase receptors (TKRs), carrying an extracellular domain for ligand binding, a transmembrane domain, and a cytoplasmic domain, that includes a tyrosine kinase domain. There are several endogenous ligands for these receptors – VEGFA/B/C/D, and placental-growth factor (PIGF), of which VEGFA and VEGFB are the most relevant. VEGFA is the one that exerts direct trophic, neuroprotective, and synaptic plasticity effects on many neuronal cells^5,6^.

VEGFA binds to and activates both VEGFR-1 and VEGFR-2, although with greater affinity for VEGFR-1^7^. The majority of the VEGFA signaling-associated effects, such as angiogenesis promotion, vascular permeability, cell migration, gene expression, neuronal survival, neuroregeneration, and synaptic plasticity facilitation are predominantly mediated by VEGFR-2 activation^8,6^. Although VEGFR-1 was identified before VEGFR-2, its role is less clear. It has high affinity for VEGFA and VEGFB, has low tyrosine kinase activity, and may act as a “decoy” receptor, preventing VEGFR-2 over-activation by competitively binding VEGFA^5,6^. VEGFB binds exclusively to VEGFR-1, having therefore less angiogenic potential than VEGFA, while also holding a neuroprotective effect^9^. VEGF is widely expressed throughout the nervous system, including neurons, astrocytes and microglia^10^, and alterations in VEGF expression have been demonstrated in both ALS patients and in mouse models. Indeed, VEGFA and VEGFR-2 mRNA expression have been shown to be decreased in the spinal cord of SOD1^G93A^ mice^11^. Furthermore, mutant SOD1 has been shown to lead to a destabilization of VEGF mRNA, resulting in reduced VEGFA expression and accelerated neurodegeneration in ALS^12^. Similarly, in spinal cord samples of ALS patients, VEGFA and VEGFR-2 expression was found to be decreased, when assessed through immunohistochemistry^13^. However, VEGF signaling in ALS is still a controversial subject since other studies either did not find decreased VEGFA levels in spinal cord tissue samples from SOD1^G93A^ mice^14^ or even reported an increase in VEGFA expression in the spinal cord of SOD1^G93A^ mice^15^. These contradictory findings, are also replicated in humans, with some studies finding either no alterations ^16,17,18,19,20^ or increases^19,21^ in VEGFA levels in the CSF, serum or plasma of ALS patients, while others report decreased VEGFA plasma^22^ and CSF levels^18^, or even an increase of VEGFA levels in both CSF and serum of ALS patients^23^.

In spite of the above-mentioned controversy, most of the evidence points towards a neuroprotective function of VEGF in ALS^6, 24,10^. Indeed, VEGF is a potent angiogenic and vascular permeability-enhancing factor, and both actions are very effective in hypoxia, therefore relevant in late stages of ALS. VEGF also exerts neuroprotective actions directly through the inhibition of apoptosis in hypoxic conditions and the stimulation of neurogenesis. Interference with the direct neuroprotective activities of VEGFR-2 renders motor neurons more susceptible to degeneration, while double transgenic SOD1^G93A^ mice overexpressing VEGFR-2 exhibit delayed disease onset, improved motor performance, and increased survival, compared with single transgenic SOD1^G93A^ mice^25^, in line with evidence that VEGFR-2 delays the degeneration of motor neurons^6, 10^.

Neuroprotective molecules, *per se*, can change along disease progression, being thus important not only the levels of the neuromodulators, but also of their receptors at different stages of the disease and under similar experimental conditions. Also, interactions between neuromodulators are known to occur. In this context, the unexpected negative influence of caffeine, a widely used non-selective adenosine receptor antagonist, in the course of the disease SOD1^G93A^ mice in this animal model^26^, has been suggested to be due to putative alterations in the levels of neurotrophic factors or their receptors, in particular of VEGF, but this possibility has not been directly addressed. Therefore, the first aim of this work was to perform a detailed molecular characterization of the adenosinergic and VEGF systems, in cortex and spinal cord, in pre-symptomatic and symptomatic SOD1^G93A^ mice, as well as in age-matched WT animals. Given the known gating effect of adenosine receptors over the actions of other neurotrophic factors^27^, and the existence of some evidence suggesting that A_2A_R activation stimulates VEGF production^28^, whereas A_2A_R gene deletion decreases the levels of another neurotrophic factor, Brain Derived Neurotrophic Factor (BDNF), the second goal of this work was to determine whether chronic administration of caffeine, would modulate VEGF, VEGFR or BDNF protein levels in the symptomatic stages of the disease.

As we have previously demonstrated for adenosine receptors in the hippocampus^29^, and for adenosinergic function in the neuromuscular junction^30,31^ of ALS mice, we were expecting that changes could already be detectable in the pre-symptomatic stage, which could even reverse in symptomatic stages. Furthermore, since degeneration at upper or lower motor neurons is not synchronic, we also hypothesized that alterations in the cortex and in spinal cord could not occur simultaneously. Lastly, we expected caffeine to affect not only some aspects of adenosinergic signaling but also of VEGF or BDNF signaling.

## Materials and Methods

### Ethical approval

All the experiments reported in this work were performed in full conformity with European Community Guideline (Directive 2010/63/EU). Experimental protocols were approved by the Instituto de Medicina Molecular João Lobo Antunes (iMM) institutional Animal Welfare Body (ORBEA-iMM), as well as by the Portuguese Competent Authority for Animal Welfare (Direcção Geral de Alimentação e Veterinária; DGAV). Throughout the planning and execution of the entire work, efforts were made to comply with the 3Rs principles of animal experimentation ethics.

### Animals

B6SJL-TgN (SOD1-G93A)1Gur/J males expressing the human G93A point mutation (glycine to alanine at residue 93) at SOD1 gene (Cu/Zn superoxide dismutase 1) and wild type (WT) B6SJLF1/J females were purchased from Jackson Laboratories (Bar Harbor, ME, USA). The breeding was performed at iMM rodent facility, and a colony was established. SOD1^G93A^ transgenic males were bred with WT females in a rotational scheme.

In this study we used SOD1^G93A^ in the pre-symptomatic (4-6 weeks old) and symptomatic stages (12-14 weeks old for adenosinergic and VEGF systems characterization, 11-16/18 weeks of age for caffeine experiments). Age-matched WT animals were used as controls. Both genders were used. Symptomatic SOD1^G93A^ mice had clear signs of paresis of the hind limbs. Pre-symptomatic mice displayed normal scores in the Rotarod test^29^. Animals were divided into cages (4-5 mice/cage) by gender, and ear tissue was collected to genotype the animals by Polymerase Chain Reaction (PCR). Animals were housed under a 12h light/12h dark cycle in a temperature-controlled room (21 ± 1°C) and 55 ± 10% humidity. Animals had ad libitum access to food and water. In each series of data, tissue collection from test and control mice were performed close in time.

### Quantitative Real-Time Polymerase Chain Reaction (qRT-PCR)

#### RNA extraction

Frozen tissue (cortex and spinal cord) was homogenized in QIAzol® Lysis Reagent (Qiagen Sciences, Maryland, USA) using a Potter-Elvehjem homogenizer with a Teflon piston. For RNA extraction the RNeasy Lipid Tissue Mini Kit (Qiagen Sciences, Maryland, USA) was used, according to manufacturer’s instructions. Total RNA obtained was eluted in RNase-free water and quantified by measuring absorbance at 260nm in a NanoDrop ND-1000 (Thermo Scientific(tm), Waltham, MA USA).

#### Quantitative RT-PCR

Complementary DNA (cDNA) was obtained from 2-5µg total RNA using the SuperScript First-Strand Synthesis System for RT-PCR (Invitrogen, Carlsbad, CA, EUA) according to manufacturer instructions. The reaction took place in a Bio-Rad C1000 Thermal Cycler (Bio-Rad, CA, USA) in the presence of 0.5mM dNTP, 50ng random hexamers, 5mM MgCl_2_, 10mM DTT, and 0.625U SuperScript II reverse transcriptase (EC 2.7.7.49, Invitrogen).

qRT-PCR was performed in a Rotorgene 6000 (Corbett Life Science, Sydney, Australia) in the presence of SYBR Green Master Mix (Applied Biosystems, Foster City, CA, USA) and 5 µM of specific primers against A_1_R, A_2A_R (see supplementary Table 1). For this reaction, Glyceraldehyde-3-phosphate dehydrogenase (GAPDH) gene was used as the internal reference. The PCR conditions included an initial denaturation for 2 min at 94°C, 50 cycles with 30s at 94°C, 90s at 60°C and 60s at 72°C, followed by a melting curve to assess the specificity of the reactions. Efficiencies were calculated from the given slopes of the LightCycler® 2.0 Instrument software (Roche Molecular Systems, Inc., Indianápolis, EUA) and fold change values were calculated according to Pfaffl equation^32^. After each qRT-PCR run, a melting point analysis was performed, to confirm primer specificity by the presence of a unique peak in the melting curve^33^. Results are expressed as A_1_R or A_2A_R mRNA expression/GAPDH.

#### Radioligand saturation binding assay

The radioligand-binding experiments were performed as previously descry bed^34^ with membrane fractions obtained from cortex and spinal cord. To quantify A_1_R density in the samples, 60-100 µg protein/well of cortex and 40-60 µg protein/well of spinal cord were incubated at RT for 2 hours with the selective high affinity A_1_R antagonist [^3^H]DPCPX ([^3^H]-8-Cyclopentyl-1,3-dipropylxanyine; American Radiolabeled Chemicals, Inc., St. Louis, MO, EUA) and 4 U/mL of Adenosine Deaminase (ADA; Merck, Darmstadt, Germany), to break down endogenous adenosine that might bind A_1_R), in a solution containing 50 mM Tris, 2 mM MgCl_2_.6H_2_O, pH 7.4 (Tris/Mg^2+^ buffer). Non-specific binding was measured in the presence of 2 µM of Xanthine Amine Congener (XAC, Tocris Bioscience, Bristol, United Kingdom), and the resulting values were subtracted from those obtained with [^3^H]DPCPX^35^ to determine the specific binding and normalized for protein concentration. Results are expressed as femtomole per mg of total protein (fmol/mg).

#### Western Blot

Dissected cortex and spinal cords samples were homogenized in Ristocetin Induced Platelet Agglutination buffer (RIPA; 50 mM Tris pH 8.0, 1 mM EDTA, 150 mM NaCl, 1% NP-40, 10% glycerol, 1% SDS) supplemented with a protease inhibitor cocktail (Complete Mini-EDTA free from Roche, Penzberg, Germany), and then centrifuged. Protein was quantified using a DC(tm) Protein Assay kit (Bio-Rad Laboratories, CA, USA). 200 µg of protein extracts were loaded and separated on a 10% SDS-PAGE gels and transferred onto PVDF membranes (Bio-Rad). Membranes were blocked with a 3% BSA in TBS-T (Tris-Buffered Saline with Tween-20 containing in mM: Tris base 20; NaCl 137 and 0.1% Tween-20) during 1h at RT, to avoid non-specific binding, and probed overnight at 4°C with the primary antibodies: rabbit-raised anti-A_1_R (1:500, sc-28,995, Santa Cruz Biotechnology, Dallas, Texas, USA)^36^, mouse-raised anti-A_2A_R (1:1500, Cat 05–717, Millipore, Burlington, MA, USA)^29,36^ and mouse raised anti-GAPDH (1:5000, Ref: AM4300, Invitrogen-Thermo Scientific, Waltham, MA, USA) as a loading control. Afterwards, membranes were incubated with the HRP-conjugated secondary antibodies (1:2500-1:10000, Bio-Rad) for 1h at RT: goat anti-mouse (Cat. #172-1011) and goat anti-rabbit (Cat. #170-6515).

Chemiluminescent detection was performed with ECL Plus Western Blotting Detection Reagent (GE Healthcare, Buckinghamshire, UK) in the ChemiDoc™ XRS^+^ System from Bio-Rad. The integrated intensity of each band was calculated using computer-assisted densitometry analysis with Image J 1.52a software (National Institutes of Health, Bethesda, MD, USA) and normalized to the integrated intensity of the housekeeping gene GAPDH. Images were prepared for printing in Image Lab software 5.2.1 (software available in ChemiDoc XRS+ system, Bio-Rad). For each protein evaluated, the chemiluminescence image was merged with the colorimetric image of the molecular weight marker. Results are expressed as A_1_R or A_2A_R protein levels per GAPDH levels.

#### Enzyme-linked Immunosorbent Assay (ELISA)

Cortex and spinal cord samples were homogenized in Ristocetin Induced Platelet Agglutination (RIPA) lysis buffer (1M Tris pH 8.0, 0.5M EDTA pH 8.0, 5M NaCl, 10% NP-40, 50% Glycerol), supplemented with cOmpleteTM Mini protease inhibitor cocktail tablets (Roche, Penzberg, Germany) and centrifuged. Protein quantification was performed using the DC(tm) Protein Assay kit (Bio-Rad Laboratories, Hercules, CA, USA). VEGFA, VEGFR-1 and VEGFR-2 protein levels were quantified with the DuoSet® ELISA Development System Kit (R&D Systems, Abingdon, UK), according to manufacturer instructions. Assessment of VEGFB protein levels was performed with the Mouse VEGFB ELISA Kit (Thermo Scientific, Massachusetts, USA), also according to manufacturer instructions. BDNF protein levels after caffeine treatment, were quantified using the BDNF Emax Immuno Assay System kit (Promega, Madison, Wisconsin, USA), according to manufacturer instructions. To improve BDNF detection, samples were acidified with HCl 1M, until reaching approximately pH 2.6. After 10 to 20 minutes, NaOH was added to neutralize the samples to pH 7.6. Absorbances were read in a microplate reader (Microplate Reader TECAN Infinite M200, Männedorf, Switzerland) at 450nm and 540nm, with the latter being subtracted to the former with the goal of correcting for differences in plate background. Results are expressed as picograms per milligram of total protein (pg/mg).

#### Chronic caffeine treatment

Caffeine powder ReagentPlus® (Merck, Darmstadt, Germany) was diluted in drinking water (herein designated as vehicle throughout the text and Figures, for the sake of simplicity) and administered from the 11^th^ week of age (just before symptoms onset) until the 16-18^th^ weeks of age (onset of hindlimb paresis). Experimental design consisted of four groups defined by genotype (WT or SOD1^G93A^) and drug treatment (tap water or caffeine (0.3mg/ml)). The dose of caffeine was chosen based on previous work (Potenza et al., 2013). During the experimental period, animals had ad libitum access to water, independently of treatment group. No substantial differences between treatment groups were detected in terms of water intake (approximately 5 – 6 mL per day per mouse). After 5-7 weeks of treatment, mice were sacrificed, cortex and spinal cord samples were removed and snap frozen in liquid nitrogen. All samples were kept at −80 °C for posterior analysis.

#### Changes in animal weight

Animal body weight was monitored throughout the chronic drug administration experiments, with animals being weighed once every 2 to 3 days. Because SOD1^G93A^ tend to generally weight less than their WT counterparts, group comparisons of absolute weight were deemed inadequate. As such, changes in the body weight of each animal were calculated by subtracting the weight of that animal on the first day of drug administration (72^nd^ day of life) to its weight at each subsequent time-point, with results being expressed in grams. To facilitate comparison of treatment effects, the area under the curve (AUC) of these changes in weight was obtained.

#### Statistical analysis

Comparisons were made between all groups, with statistical significance level (α) established at p < 0.05. Data was analyzed through two-way analysis of variance (ANOVA) with Holm-Sidak correction for multiple comparisons where appropriate. Data are expressed as mean ± standard error of mean (SEM). All statistical analyses were performed using GraphPad Prism 8 software (GraphPad Software, San Diego, CA, USA).

## Results

### Adenosine receptor levels were altered in the cortex of SOD1^G93A^ mice

Analysis of cortical A_1_R mRNA expression revealed no significant main effects of either genotype (F_(1, 19)_ = 0.623, p = 0.440) or age (F_(1, 19)_ = 1.83, p = 0.193), but a significant genotype x age interaction (F_(1, 19)_ = 5.77, p = 0.03). Congruently, no significant differences were observed in cortical A_1_R mRNA expression in pre-symptomatic (WT = 0.902 ± 0.2148, pre-symptomatic SOD1^G93A^ = 1.538 ± 0.2419, p = 0.184) or symptomatic SOD1^G93A^ mice (WT = 1.112 ± 0.219, symptomatic SOD1^G93A^ = 0.790 ± 0.0816, p = 0.59), in relation to their respective age-matched WT congeners (Figure 1A).

**Figure 1.**
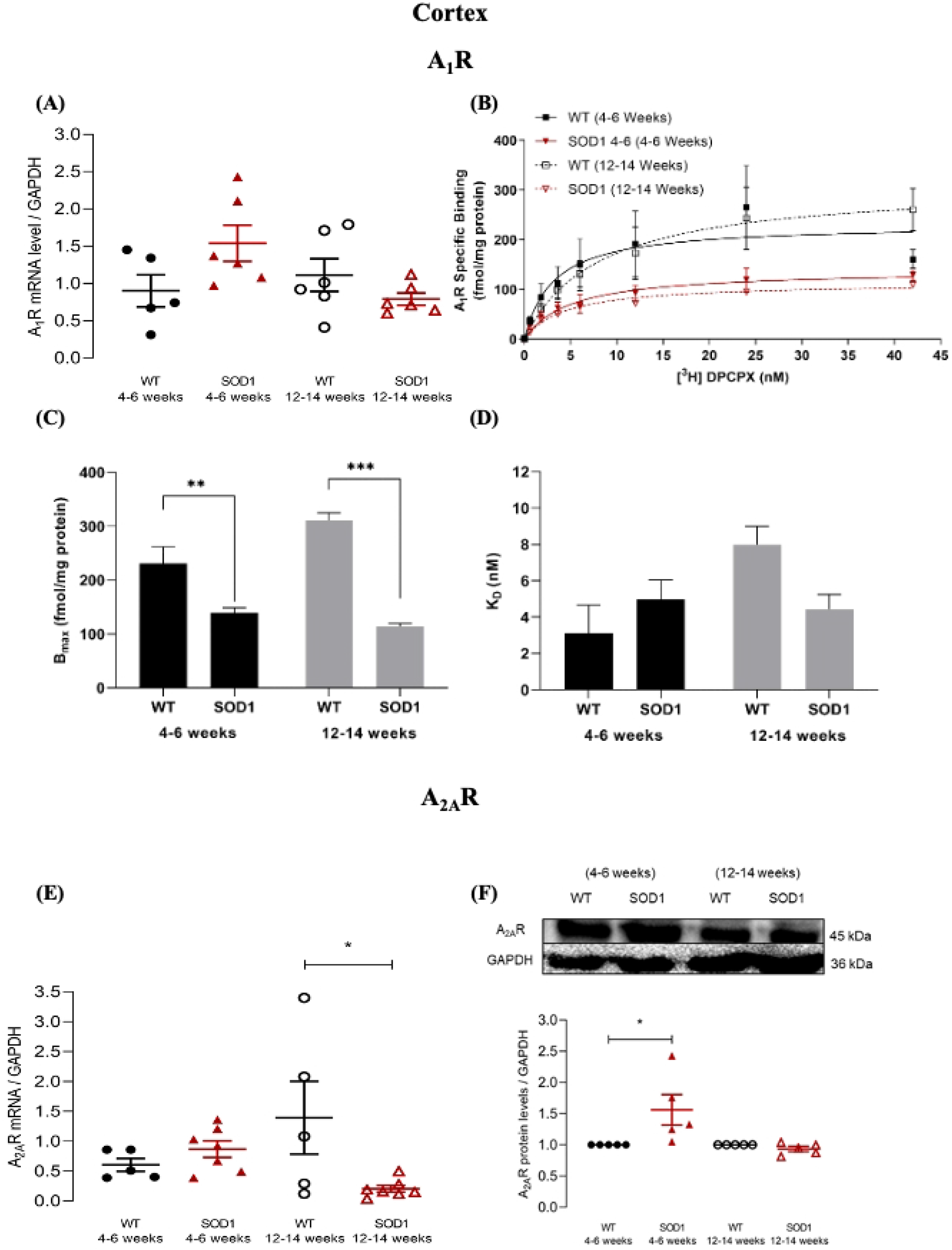
A_1_R and A_2A_R mRNA expression and protein levels in the cortex of pre-symptomatic and symptomatic SOD1^G93A^ mice. **(A)** Cortical A_1_R mRNA expression in pre-symptomatic (4-6 weeks old) and symptomatic (12-14 weeks old) SOD1^G93A^ and age-matched WT mice. The Y-axis represents the A_1_R transcript level of the normalized A_1_R/GAPDH ratio in each condition. **(B)** Saturation binding curves of A_1_R for pre-symptomatic and symptomatic SOD1^G93A^ and age-matched WT mice, in cortex. The Y-axis represents the specific binding of A_1_R expressed in fmol/mg protein plotted against increased concentrations of [^3^H] DPCPX (nM) (abscissae). **(C)** Maximum density of A_1_R (Bmax) in cortex of pre-symptomatic and symptomatic SOD1^G93A^ and age-matched WT mice. **(D)** DPCPX equilibrium dissociation constant (K_D_) expressed in nM, for pre-symptomatic and symptomatic SOD1^G93A^ and age-matched WT mice, in cortex. **(E)** Cortical A_2A_R mRNA expression in pre-symptomatic and symptomatic SOD1^G93A^ and age-matched WT mice. The Y-axis represents the A_2A_R transcript level of the normalized A_2A_R/GAPDH ratio in each condition. **(F)** Cortical A_2A_R protein levels in pre-symptomatic and symptomatic SOD1_G93A_ and age-matched WT mice. The Y-axis represents the A_2A_R/GAPDH ratios normalized to age-matched WT samples. Upper panels show representative immunoblots for each condition. Data are expressed as mean ±SEM (n=4-7 for all conditions); *p < 0.05, **p ≤ 0.01, ***p ≤ 0.001, two-way ANOVA with Holm-Sidak correction for multiple comparisons.

Interestingly, and in contrast with the mRNA results, analysis of cortical A_1_R density revealed a significant main effect of genotype (F_(1, 16)_ = 87.4, p < 0.001), but not a significant main effect of age (F_(1, 16)_ = 3.09, p = 0.098). Moreover, a significant genotype x age interaction (F(1, 16) = 11.7, p = 0.004) was observed. Post-hoc pair-wise tests showed a significant decrease of the maximum number of specific A_1_R binding sites (B_max_) in SOD1^G93A^ mice, in both the pre-symptomatic (WT = 231.1 ± 30.82, pre-symptomatic SOD1^G93A^ = 139.3 ± 9.323, p = 0.001), and symptomatic stages (WT = 311.1 ± 13.93, symptomatic SOD1^G93A^ = 113.7 ± 6.258, p < 0.001) (Figures 1B and C). When the cortical A_1_R affinity was assessed, no significant main effects either of genotype (F_(1, 16)_ = 0.523, p = 0.480) or age (F_(1, 16)_ = 3.48, p = 0.081) were found, but a significant main effect of genotype x age interaction was found (F_(1, 16)_ = 5.42, p = 0.033). No K_D_ differences were observed between the pre-symptomatic (WT = 3.101 ± 1.55, pre-symptomatic SOD1^G93A^ = 4.97 ± 1.072; p = 0.59) or symptomatic stages (WT = 7.983 ± 1.012, symptomatic SOD1^G93A^ = 4.431 ± 0.8213, p = 0.239) (Figure 1D).

With regards to cortical A_2A_R mRNA expression, while analysis revealed no significant main effects of either genotype (F_(1, 20)_ = 2.90, p = 0.104) or age (F_(1, 20)_ = 0.056, p = 0.815), a significant genotype x age interaction (F_(1, 20)_ = 7.15, p = 0.015) was observed. Post-hoc tests found that, while during the pre-symptomatic stage no difference in cortical A_2A_R mRNA was observed (WT = 0.604 ± 0.107, pre-symptomatic SOD1^G93A^ = 0.867 ± 0.138, p = 0.526), in the symptomatic stage a significant decrease became evident (WT = 1.394 ± 0.6097, SOD1^G93A^ = 0.2057 ± 0.0561, p = 0.034) (Figure 1E). Analysis of cortical A_2A_R protein levels revealed no significant main effect of genotype (F_(1, 16)_ = 3.99, p = 0.063), but a significant effect was found for age (F_(1, 16)_ = 6.38, p = 0.022), in addition to a significant genotype x age interaction (F_(1, 16)_ = 6.38, p = 0.022). Post-hoc pair-wise tests revealed a significant increase of A_2A_R protein levels in the cortex of pre-symptomatic SOD1^G93A^ mice (pre-symptomatic SOD1^G93A^ = 1.562 ± 0.245, p = 0.011), but not in the symptomatic stage (SOD1^G93A^ = 0.934 ± 0.041, p = 0.997) (Figure 1F).

Summarizing, in the cortex of SOD1^G93A^ mice, A_1_R levels were significantly decreased both in pre-symptomatic as well as in the symptomatic states of the disease, whereas A_2A_R protein levels were increased in pre-symptomatic stages. A_2A_R mRNA levels were decreased in symptomatic SOD1^G93A^ mice, which may lead to the normalization of the A_2A_R levels in the symptomatic stages of the disease.

### Adenosine receptors levels were altered in the spinal cord of SOD1^G93A^ mice

Concerning spinal cord A_1_R mRNA expression, while no significant genotype main effect was observed (F_(1, 20)_ = 1.83, p = 0.191), a significant main effect of age (F_(1, 20)_ = 5.61, p = 0.028), and a significant genotype x age interaction (F_(1, 20)_ = 7.28, p = 0.014) were detected. Post-hoc comparisons revealed that, in comparison to age-match WT animals, pre-symptomatic SOD1^G93A^ animals showed increased spinal cord A_1_R mRNA expression (WT = 0.9250 ± 0.1384, pre-symptomatic SOD1^G93A^ = 2.137 ± 0.557, p = 0.047). As the disease progressed these differences in mRNA expression seemed to be reverted, as evidenced by both a significant difference between SOD1^G93A^ mice in the pre-symptomatic and symptomatic stages (symptomatic SOD1^G93A^ = 0.622 ± 0,135, p = 0.011) and the existence of no significant differences between symptomatic SOD1^G93A^ mice and age-match WT controls (WT = 1.023 ± 0.1, p = 0.729) (Figure 2A).

**Figure 2.**
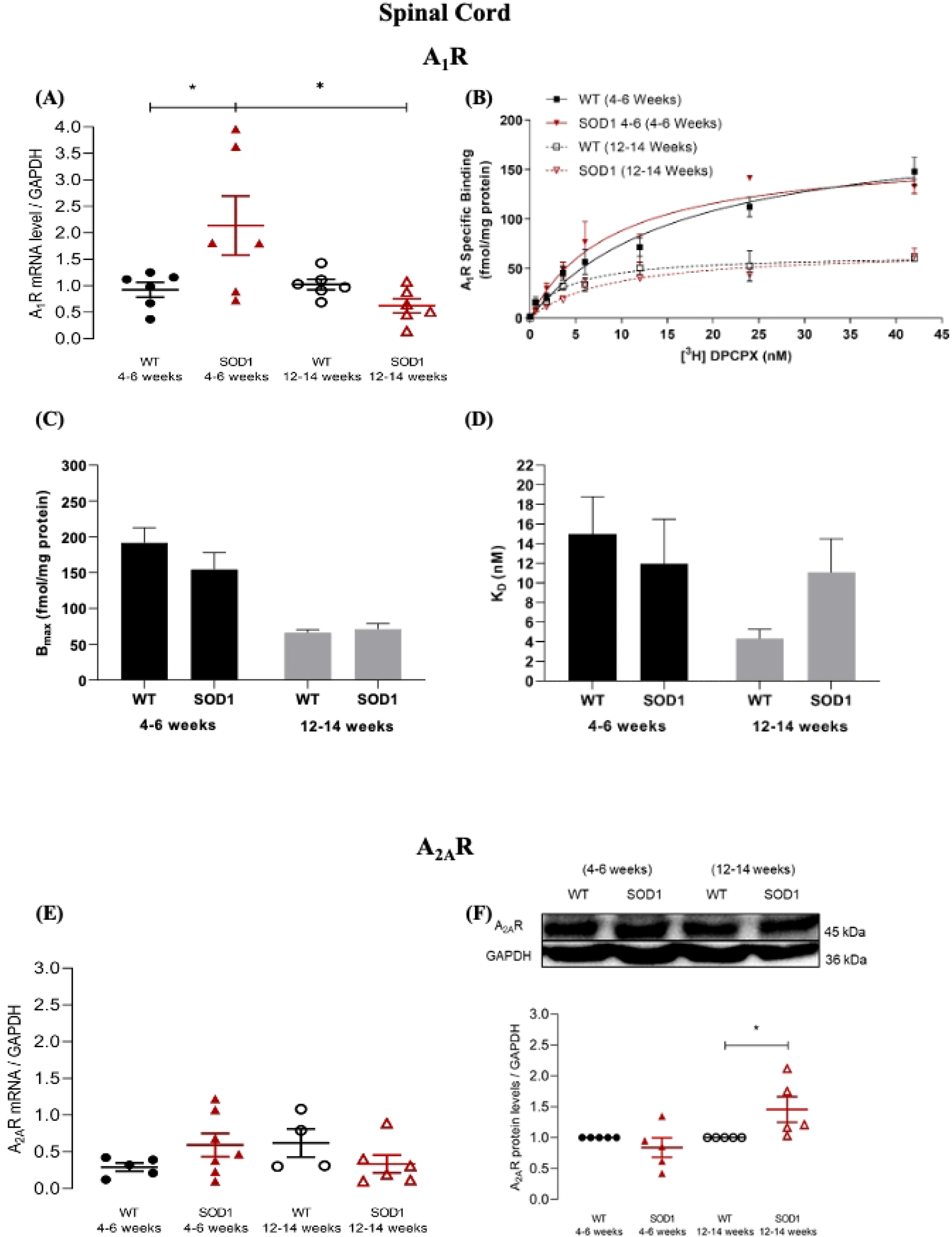
A_1_R and A_2A_R mRNA expression and protein levels in the spinal cord of pre-symptomatic and symptomatic SOD1^G93A^ mice. **(A)** A_1_R mRNA expression in spinal Cord of pre-symptomatic (4-6 weeks old) and symptomatic (12-14 weeks old) SOD1^G93A^ and age-matched WT mice. The Y-axis represents the A_1_R transcript level of the normalized A_1_R/GAPDH ratio in each condition. **(B)** Saturation curve of A_1_R for pre-symptomatic and symptomatic SOD1^G93A^ and age-matched WT mice, in spinal cord. The Y-axis represents the specific binding of A_1_R expressed in fmol/mg protein plotted against increased concentrations of [^3^H] DPCPX (nM) (abscissae). **(C)** Maximum density of A_1_R (Bmax) in spinal cord of pre-symptomatic and symptomatic SOD1^G93A^ and age-matched WT mice. **(D)** DPCPX equilibrium dissociation constant (K_D_) expressed in nM, for pre-symptomatic and symptomatic SOD1^G93A^ and age-matched WT mice, in spinal cord. **(E)** A_2A_R mRNA expression in spinal cord of pre-symptomatic and symptomatic SOD1^G93A^ and age-matched WT mice. The Y-axis represents the A_2A_R transcript level of the normalized A_2A_R/GAPDH ratio in each condition **(F)** A_2A_R protein levels in spinal cord of pre-symptomatic and symptomatic SOD1^G93A^ and age-matched WT mice. The Y-axis represents the A_2A_R/GAPDH ratios normalized to age-matched WT samples. Upper panels show representative immunoblots for each condition. Data are expressed as mean ±SEM (n=4-7 for all conditions); *p < 0.05, two-way ANOVA with Holm-Sidak correction for multiple comparisons.

Regarding A_1_R levels in spinal cord, no significant main effect of genotype (F(1, 22) = 0.725, p = 0.404), but a significant main effect of age was found (F(1, 22) = 38.1, p < 0.001). No significant genotype x age interaction (F(1, 22) = 1.28, p = 0.270) was detected. Post-hoc pair-wise tests showed no differences in the spinal cord of mice when compared with age-matched WT (pre-symptomatic: WT = 198.6 ± 21.35, SOD1^G93A^ = 162.2 ± 17.68, p = 0.225; symptomatic: WT = 64.52 ± 4.265, SOD1^G93A^ = 69.65 ± 6.437, p = 0.861) (Figure 2B and C).

No differences were found in A_1_R KD values in either pre-symptomatic (WT = 14.97 ± 3.828, SOD1^G93A^ = 11.96 ± 4.554; p = 0.867; Fig. 3.4f) or symptomatic SOD1^G93A^ animals (WT = 4.322 ± 0.924, SOD1^G93A^ = 11.05 ± 3.467, p = 0.792, in relation to age-matched WT mice (Figure 2D).

With regards to spinal cord A_2A_R mRNA expression, no evidence for a significant main effect either of genotype (F_(1, 18)_ = 0.003, p = 0.958), or age (F_(1, 17)_ = 0.058, p = 0.813) was achieved, but a tendency towards a significant genotype x age interaction (F_(1, 17)_ = 4.07, p = 0.059) was observed. Pair-wise post-hoc comparisons found no significant differences either at the pre-symptomatic (WT = 0.292 ± 0.056, pre-symptomatic SOD1^G93A^ = 0.593 ± 0.159, p = 0.597) or symptomatic disease stages (WT = 0.620 ± 0.191, symptomatic SOD1^G93A^ = 0.335 ± 0.120, p = 0.597) (Figure 2E).

Concerning spinal cord A_2A_R protein levels, while no significant main effect of genotype (F_(1, 16)_ = 1.29, p = 0.273) was observed, a significant main effect was reached for age (F_(1, 16)_ = 5.68, p =.030), in addition to a significant genotype x age interaction (F_(1, 16)_ = 5.68, p = 0.030). Post-hoc pair-wise tests revealed no differences in pre-symptomatic stage (pre-symptomatic SOD1^G93A^ = 0.838 ± 0.157, p = 0.391), and a significant increase at the symptomatic stage (symptomatic SOD1^G93A^ = 1.455 ± 0.206, p = 0.048) (Figure 2F).

Summarizing data obtained with the spinal cord, A_1_R protein levels were unaltered in SOD1^G93A^ mice as compared with age-matched WT mice. There was, however a moderate but significant increase in A_1_R mRNA levels in the pre-symptomatic mice, but that does not reflect into changes in protein levels. A_2A_R protein levels were significantly increased only in the symptomatic SOD1^G93A^ mice.

Altogether, data point towards an earlier alteration in the cortex than in spinal cord. Whenever occurring, the direction of the change is for a decrease in A_1_R and an increase in A_2A_R.

### VEGF system were altered in the cortex of SOD1^G93A^ mice

Concerning VEGFA protein levels in the cortex, analysis revealed no significant main effect of genotype (F_(1, 16)_ = 0.072, p = 0.792), a significant main effect of age (F_(1, 16)_ = 11.3, p = 0.004), and no significant effect of genotype x age interaction (F_(1, 16)_ = 0.058, p = 0.813). Pair-wise post-hoc comparisons indicated no differences either in pre-symptomatic (WT = 6.389 ± 0.581, pre-symptomatic SOD1^G93A^ = 5.985 ± 1.039, p = 0.936) or symptomatic SOD1^G93A^ animals (WT = 3.525 ± 0,764, symptomatic SOD1^G93A^ = 3.504 ± 0.689, p = 0.983) in relation to their age-matched WT congeners (Figure 3A).

**Figure 3.**
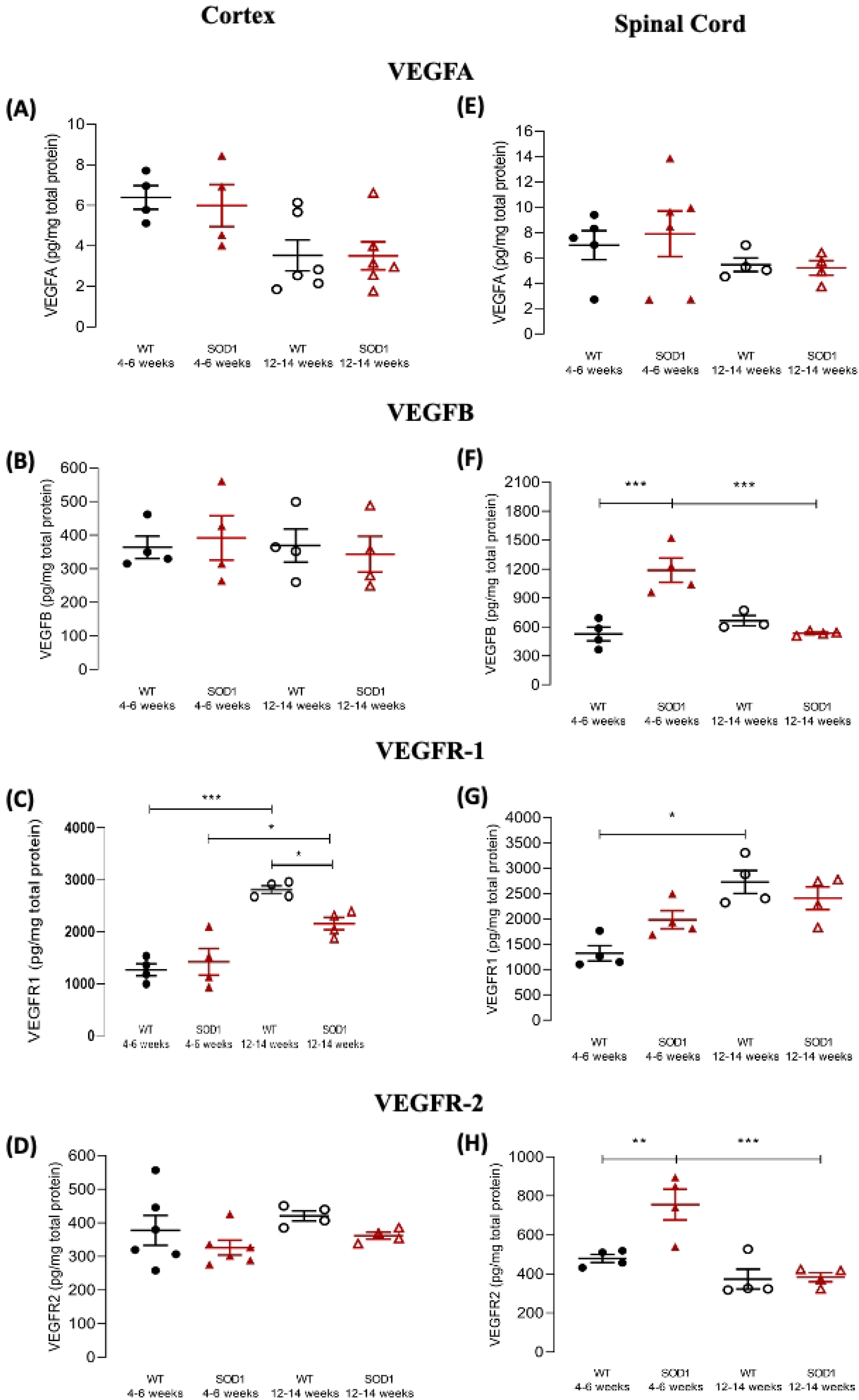
VEGF and VEGF receptor protein levels in the cortex and spinal cord of pre-symptomatic and symptomatic SOD1G93A mice. **(A-D)** Data from cortical samples; **(E-H)** data from spinal cord samples. VEGFA **(A, E)**, VEGFB **(B, F)**, VEGFR-1 **(C, G)** or VEGFR-2 **(D, H)** protein levels, determined by ELISA, in samples from pre-symptomatic (4-6 weeks old) or symptomatic (12-14 weeks old) SOD1G93A and age-matched WT mice, as indicated below each data set. The Y-axis represents the ligand or receptor levels expressed in pg/mg of total protein in each condition. Data are expressed as mean ± SEM (n = 3-7); *p < 0.05, **p ≤ 0.01, ***p ≤ 0.001, two-way ANOVA with Holm-Sidak correction for multiple comparisons.

For cortical VEGFB protein levels, no evidence was found for a significant main effect of either genotype (F_(1, 12)_ = 0.0003, p = 0.986), or age (F_(1, 12)_ = 0.177, p = 0.682), nor for a significant genotype x age interaction (F_(1, 12)_ = 0.264, p = 0.617). Congruently, pair-wise post-hoc comparisons revealed no differences in either the pre-symptomatic (WT = 364.2 ± 33.24, pre-symptomatic SOD1^G93A^ = 391.7 ± 65.85, p = 0.998) or the symptomatic disease stages (WT = 369.0 ± 49.29, symptomatic SOD1^G93A^ = 343.4 ± 53.22, p = 0.998) (Figure 3B).

Analysis of cortical VEGFR-1 protein levels showed no significant main effect of genotype (F_(1, 12)_ = 2.50, p = 0.140), but a significant main effect of age (F_(1, 12)_ = 52.3, p < 0.001), as well as a significant effect of genotype x age interaction (F_(1, 12)_ = 6.60, p = 0.025). Pair-wise post-hoc comparisons revealed no differences between pre-symptomatic mice and age-matched WT mice (WT = 1270 ± 115.0, pre-symptomatic SOD1^G93A^ = 1425 ± 256.7, p = 0.498), and a slight decrease in symptomatic mice when compared with age-matched WT animals (WT = 2810 ± 73.73, symptomatic SOD1^G93A^ = 2158 ± 119.9, p = 0.025). Significant increases were found between WT animals of different ages (p < 0.001), as well as between pre-symptomatic and symptomatic SOD1^G93A^ mice (p = 0.019) (Figure 3C).

With regards to VEGFR-2, analysis of cortical VEGFR-2 protein levels showed no significant main effect of genotype (F_(1, 16)_ = 3.01, p = 0.102), nor of age (F_(1, 16)_ = 1.53, p = 0.234). Likewise, no evidence for a significant genotype x age interaction was observed (F_(1, 16)_ = 0.0127, p = 0.912). Congruently, pair-wise post-hoc comparisons revealed no differences between pre-symptomatic (WT = 377.8 ± 44.48, pre-symptomatic SOD1^G93A^ = 326.1 ± 21.95, p = 0.707) or symptomatic SOD1^G93A^ mice (WT = 420.7 ± 15.04, symptomatic SOD1^G93A^ = 361.9 ± 10.27, p = 0.707) in relation to their respective age-matched WT controls (Figure 3D).

In summary, in the cortex of SOD1^G93A^ mice the protein levels of VEGFA and of VEGFB were not significantly altered as compared with age-matched WT mice. In relation to the receptors, no significant changes were detected in VEGFR-2 protein levels in SOD1^G93A^ mice as compared with WT. For VEGFR-1, the age-dependent increase in protein level was moderate in SOD1^G93A^ mice and accentuated in WT mice, turning theVEGFR-1 protein levels significantly lower in symptomatic mice as compared with age-matched WT mice.

### VEGF system was altered in the spinal cord of SOD1^G93A^ mice

Analysis of spinal cord VEGFA protein levels revealed no significant main effects of genotype (F_(1, 15)_ = 0.053, p = 0.821), or age (F_(1, 15)_ = 2.38, p = 0.144), nor the existence of a significant effect of genotype x age interaction (F_(1, 15)_ = 0.175, p = 0.681). Congruently, pair-wise post-hoc comparisons found no differences between SOD1^G93A^ mice either at the pre-symptomatic (WT = 7.005 ± 1.144, pre-symptomatic SOD1^G93A^ = 7.896 ± 1.798, p = 0.860) or symptomatic stages (WT = 5.461 ± 0.537, symptomatic SOD1^G93A^ = 5.202 ± 0.574, p = 0.903) when compared with age-matched WT mice, nor between the two disease stages (p = 0.690) (Figure 3E).

Regarding VEGFB protein levels, a significant main effect of genotype (F_(1, 11)_ = 10.7, p = 0.007), a significant main effect of age (F_(1, 11)_ = 10.1, p = 0.009), and a significant main effect of genotype x age interaction (F_(1, 11)_ = 23.8, p < 0.001) was detected. Pair-wise post-hoc comparisons revealed a significant increase in VEGFB protein levels in the spinal cord of pre-symptomatic SOD1^G93A^ mice, comparing with age-matched wild type (WT = 526.7 ± 71.10, pre-symptomatic SOD1^G93A^ = 1188 ± 125.0, p < 0.001), but no such differences were observed in symptomatic mice when compared with age-matched WT mice (WT = 664.8 ± 53.27, symptomatic SOD1^G93A^ = 534.5 ± 11.29, p = 0.612). Therefore, a significant (p < 0.001) decrease in VEGFB protein levels was detected, when comparing pre-symptomatic and symptomatic SOD1^G93A^ mice, while such difference was not observed for WT animals (p = 0.612) (Figure 3F).

Assessment of spinal cord VEGFR-1 protein levels showed a significant main effect of genotype (F_(1, 12)_ = 0.742, p = 0.406), a significant main effect of age (F_(1, 12)_ = 21.4, p < 0.001), and a significant genotype x age interaction (F_(1, 12)_ = 6.1, p = 0.029). Pair-wise post-hoc comparisons revealed no differences in the spinal cord of pre-symptomatic mice when compared with age-matched WT (WT = 1320 ± 152.5, pre-symptomatic SOD1^G93A^ = 1982 ± 178.3, p = .103), nor any differences at the symptomatic stage (WT = 2728 ± 228.1, symptomatic SOD1^G93A^ = 2407 ± 223.1, p = 0.286). However, a significant increase was found in older vs. younger WT animals (p = 0.002), while no such difference was observed for SOD1^G93A^ mice (p = 0.286) (Figure 3G).

With regards to VEGFR-2 protein levels, a significant main effect of genotype (F_(1, 12)_ = 8.33, p = 0.014), a significant main effect of age (F_(1, 12)_ = 23.2, p < 0.001) and a significant genotype x age interaction (F_(1, 12)_ = 7.19, p = 0.020). Pair-wise post-hoc comparisons revealed a significant increase of VEGF-R2 levels in pre-symptomatic SOD1^G93A^ mice (WT = 479.7 ± 20.90, pre-symptomatic SOD1^G93A^ = 755.7 ± 78.96, p = 0.008), while no such difference was observed in symptomatic animals (WT = 373.8 ± 51.11, symptomatic SOD1^G93A^ = 383.9 ± 23.41, p = 0.888). Therefore, comparison between pre-symptomatic and symptomatic SOD1^G93A^ animals revealed a statistically significant decrease in VEGF-R2 from the pre-symptomatic to the symptomatic stage (p < 0.001) while no such change was observed in WT animals (Figure 3H).

Summarizing the data obtained with spinal cord samples, we detected an increase in VEGFB and VEGFR-2 levels in the pre-symptomatic as compared with age-matched WT mice, which was no longer evident in the symptomatic mice. This led to significant decreases in the levels of those proteins in the symptomatic vs the pre-symptomatic disease stage.

### Caffeine treatment did not affect the weight-gain SOD1^G93A^ mice

Caffeine was orally administered from the 11^th^ week of age (just before onset of symptoms) until the 16^th^-18^th^ weeks of age (onset of hindlimb paresis). During the treatment period, animal weight was recorded three times per week, and weight change was plotted taking as zero the weight immediately before caffeine administration (Post-Natal Day (PND) 72) (Figure 4A). To assess putative changes in body weight caused by caffeine in WT and in SOD SOD1^G93A^ mice, we calculated the area under the curve (AUC) of the average change in body weight in relation to the weight on the first day of drug administration. Two-way ANOVA of the AUC revealed a significant main effect of genotype (F_(1, 36)_ = 282, p < 0.001) and a significant main effect of treatment (F_(1, 36)_ = 9.65, p = 0.004). However, the genotype x treatment interaction (F_(1, 36)_ = 0.115, p = 0.737) was not significant. Pair-wise post-hoc comparisons revealed a significant decrease in weight-gain in vehicle-treated SOD1^G93A^ mice in comparison to vehicle-treated WT mice (WT-VEH = 71.20 ± 2.011; SOD-VEH =12.90 ± 3.033, p <.001), and a similar significant decrease in weight gain in caffeine-treated SOD1^G93A^ animals in relation to their caffeine-treated WT counterparts (WT-CAF = 83.40 ± 4.743, SOD-CAF = 22.70 ±3.795, p < 0.001). In WT animals, caffeine treatment led to a small but statistically significant increase in weight gain, as compared to vehicle treatment (p = 0.039). Similarly, a trend towards an increase in weight gain was observed in caffeine-treated SOD1^G93A^ animals in relation to their vehicle-treated SOD1^G93A^ congeners, but it did not reach statistically significance (p = 0.058).

**Figure 4.**
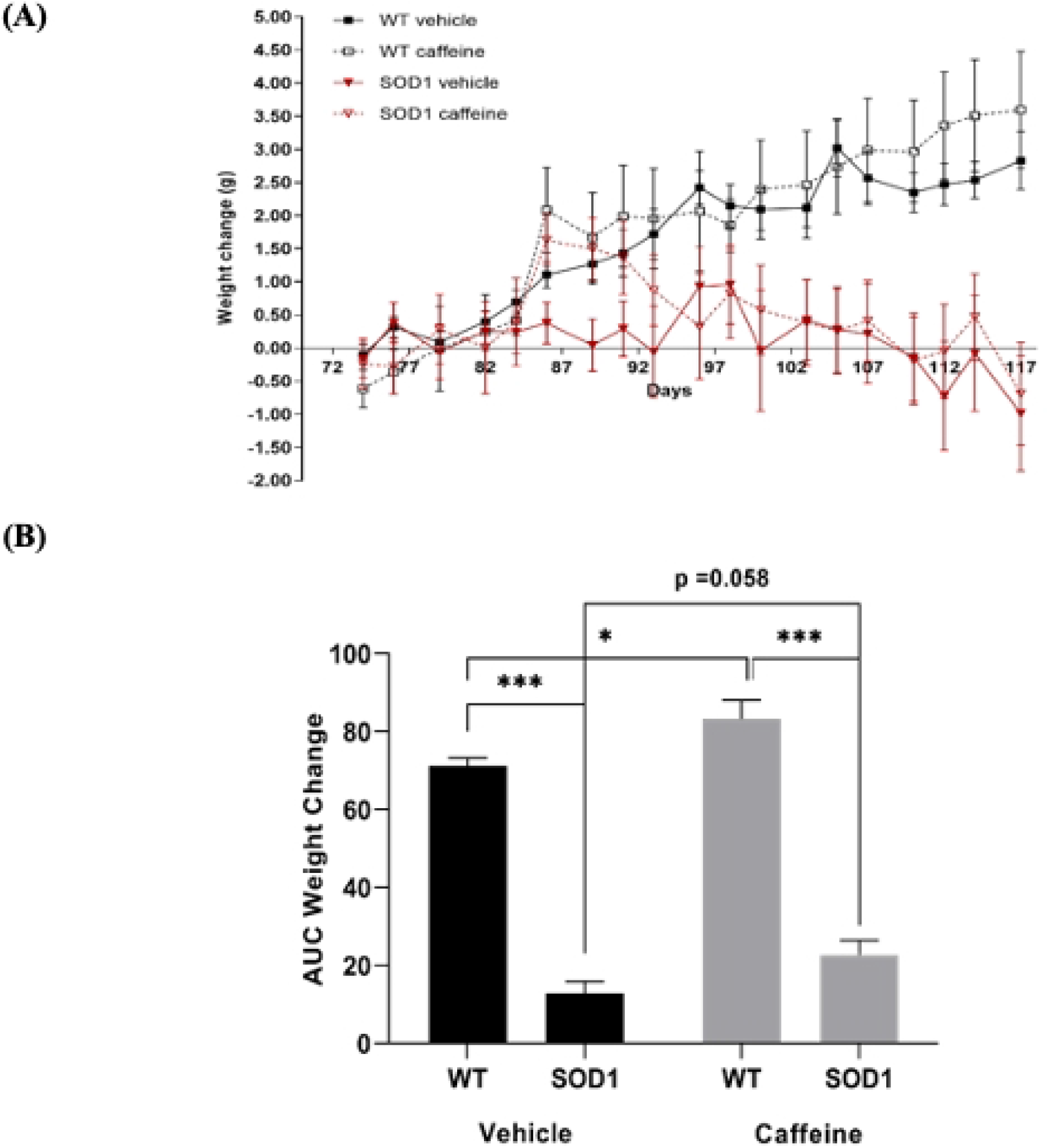
Effect of chronic caffeine treatment on animal weight gain over time. **(A)** The Y-axis represents the weight change caffeine treated/non-treated WT or SOD1^G93A^ animals in relation to the first day of treatment. The X-axis represents the days of treatment. **(B)** Area under the curve (AUC) of weight change for caffeine treated/non-treated WT or SOD1^G93A^ animals. Data are expressed as mean ± SEM (n = 6-10); *p < 0.05; ***p ≤ 0.001, two-way ANOVA with Holm-Sidak correction for multiple comparisons.

Visual inspection of the mice showed a trend to symptoms aggravation in the SOD1^G93A^ mice treated with caffeine, in accordance with what has been previously reported by Potenza et al (2013). Accordingly, caffeine treated SOD1^G93A^ mice reached the defined humane endpoint (onset of hindlimb paresis) slightly earlier than vehicle treated SOD1^G93A^ mice. To avoid bias in the molecular analysis, whenever a SOD1^G93A^ mouse reached the humane endpoint, one mouse of all the other groups (caffeine or vehicle treated SOD1^G93A^ mouse, caffeine or vehicle treated WT mouse) were also euthanized. Objective analysis of the influence of caffeine on the lifespan of the animals cannot, therefore, be done, but it was not under the scope of our work since it has been already reported by others (Potenza et al., 2013).

### Caffeine treatment did not affect the VEGF system in the cortex of SOD1^G93A^ mice

While assessing the influence of caffeine upon the VEGFA protein levels protein levels in the cerebral cortex of SOD1^G93A^ mice and WT mice, a two-way ANOVA taking the genotype (WT or SOD1^G93A^) and treatment (vehicle or caffeine) as the factors revealed a significant main effect of genotype (F(1, 14) = 12.0, p = 0.004), but no significant main effect of treatment (F(1, 14) = 2.66, p = 0.125). However, a significant genotype x treatment interaction (F(1, 14) = 12.5, p = 0.003) was observed. Pair-wise post-hoc comparisons revealed a significant decrease in VEGFA protein levels in vehicle-treated SOD1^G93A^ mice when compared with vehicle-treated WT animals (WT-VEH = 3.615 ± 0.434, SOD1-VEH = 1.492 ± 0.280, p = 0.002). This observation, when compared with the one shown in Figure 3A suggests a decline in VEGF levels in late stage (16-18 weeks old) SOD1^G93A^ mice (Figure 5, vehicle treated SOD1^G93A^ mice significantly different from treatment and age-matched WT mice) that was absent in early symptomatic (12-14 weeks old mice, no differences between SOD1^G93A^ and WT mice). No differences were found between WT and SOD1^G93A^ mice treated with caffeine (WT-CAF= 2.047 ± 0.324, p = 0.956), indicating an absence of effect of caffeine upon VEGFA protein levels (Figure 5A).

**Figure 5.**
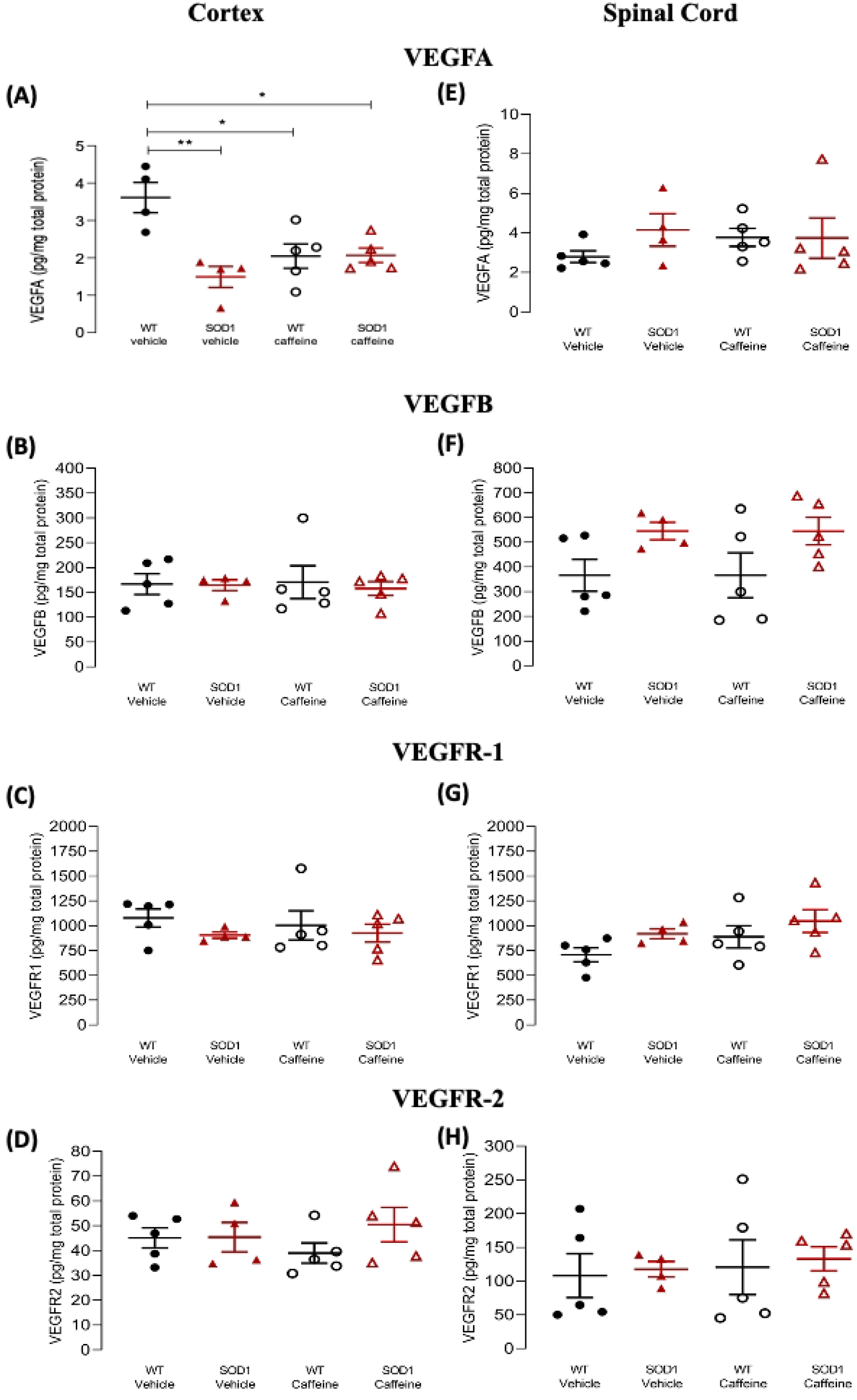
VEGF and VEGF receptor protein levels in the cortex and spinal cord of late stage SOD1^G93A^ and WT mice treated/non-treated with caffeine. **(A-D)** Data from cortical samples; **(E-H)** Data from spinal cord samples. VEGFA **(A, E)**, VEGFB **(B, F)**, VEGFR-1 **(C-G)** and VEGFR-2 **(D-H)** protein levels in samples from late stages SOD1^G93A^ mice (16-18 weeks old) and age-matched WT mice, treated/not treated with caffeine (start at the age of 11 weeks), as indicated in each panel below each data set. Caffeine administration in the drinking water (see methods) started when animals were 11 weeks old. The Y-axis represents the ligands or receptor protein levels, quantified by ELISA, in pg/mg of total protein. Data are expressed as mean ± SEM (n = 4-6); *p < 0.05, **p ≤ 0.01, two-way ANOVA with Holm-Sidak correction for multiple comparisons.

Considering cortical VEGFB protein levels, a two-way ANOVA revealed no significant main effects either of genotype (F(1, 15) = 0.111, p = 0.744), or treatment (F(1, 15) = 0.00421, p = 0.949), nor a significant genotype x treatment interaction (F(1, 15) = 0.0580, p = 0.813). In line with this, pair-wise post-hoc comparisons revealed no differences between vehicle-treated WT and SOD1^G93A^ animals (WT-VEH = 166.8 ± 20.91, SOD-VEH = 164.7 ± 10.79, p > 0.999), nor between caffeine-treated WT and SOD1^G93A^ mice (WT-CAF= 170.7 ± 33.07, SOD-CAF = 157.8 ± 13.94, p = 0.999) (Figure 5B). Cortical VEGFR-1 protein levels (Fig. 4.4e) were also not affected by caffeine treatment. A two-way ANOVA showed no significant main effects of genotype (F(1, 15) = 1.44, p = 0.248), or treatment (F(1, 15) = 0.064, p = 0.803), nor a significant genotype x treatment interaction (F(1, 15) = 0.210 p = 0.653). Congruently, post-hoc comparisons revealed no significant differences between WT and SOD1^G93A^ animals treated with the vehicle solution (WT-VEH = 1078 ± 90.79, SOD-VEH = 905.1 ± 30.64, p = 0.851), nor between caffeine-treated WT and SOD1^G93A^ animals (WT-CAF= 1004 ± 146.9, SOD-CAF = 926.5 ± 89.92, p = 0.949) (Figure 5C).

Similarly, caffeine treatment did not alter VEGFR-2 protein levels in the cerebral cortex. A two-way ANOVA, revealed no significant main effect for genotype (F(1, 15) = 1.20, p = 0.291), or treatment (F(1, 15) = 0.009, p = 0.926). Congruently the genotype x treatment interaction was also non-significant (F(1, 15) = 1.09, p = 0.313). Pair-wise post-hoc comparisons revealed no differences either between vehicle-treated WT and SOD1^G93A^ animals (WT-VEH = 45.14 ± 4.019, SOD-VEH = 45.41 ± 5.938, p = 0.974), nor between caffeine-treated WT and SOD1^G93A^ animals (WT-CAF = 39.02 ± 4.077, SOD-CAF = 50.51 ± 6.925, p = 0.595) (Figure 5D).

### Caffeine treatment did not affect VEGF or VEGFR protein levels in the spinal cord of SOD1^G93A^ mice

With regards to VEGFA protein levels, two-way ANOVA revealed a non-significant main effect of genotype (F(1, 15) = 0.890, p = 0.360), a non-significant main effect of treatment (F(1, 15) = 0.165, p = 0.690), and also a non-significant genotype x treatment interaction (F(1, 15) = 0.977, p = 0.339). Pair-wise post-hoc comparisons revealed no differences between vehicle-treated WT and SOD1^G93A^ mice (WT-VEH = 2.798 ± 0.296, SOD1-VEH = 4.151 ± 0.820, p = 0.746), nor between caffeine-treated WT and SOD1^G93A^ mice (WT-CAF = 3.774 ± 0.451, SOD-CAF= 3.743 ± 1.018, p = 0.974) (Figure 5E).

Concerning spinal cord VEGFB protein levels, two-way ANOVA found a significant main effect of genotype (F(1, 15) = 6.95, p = 0.019), but no significant main effect of treatment (F(1, 15) = 4.80×10-7, p > 0.999), nor a significant genotype x treatment interaction (F(1, 15) = 3.18×10-5, p = 0.996). Pair-wise post-hoc comparisons revealed no differences between vehicle-treated WT and SOD1^G93A^ mice (WT-VEH= 366 ± 64.68, SOD-VEH = 545.6 ± 35.13, p = 0.369), nor between WT and SOD1^G93A^ animals treated with caffeine (WT-CAF = 366.5 ± 90.97, SOD-CAF = 545.3 ± 55.72, p = 0.369) (Figure 5F).

Regarding spinal cord VEGFR-1 protein levels, two-way ANOVA found a trend toward significant main effect of genotype (F(1, 15) = 3.7, p = 0.072), but no significant main effect of treatment (F(1, 15) = 2.61, p = 0.127), nor a significant genotype x treatment interaction (F(1, 15) = 0.075, p = 0.788). Congruently, pair-wise post-hoc comparisons revealed no significant differences between vehicle-treated WT and SOD1^G93A^ groups (WT-VEH= 707.9 ± 70.19, SOD-VEH= 919.3 ± 50.34, p = 0.557), nor between the caffeine-treated WT and SOD1^G93A^ animals (WT-CAF= 889 ± 112.5, SOD-CAF= 1048 ± 115.2, p = 0.570) (Figure 5G).

Lastly, analysis of spinal cord VEGFR-2 protein levels, two-way ANOVA found no significant main effects either for genotype (F(1, 15) = 0.135, p = 0.719), or treatment (F(1, 15) = 0.224, p = 0.643), nor a significant genotype x treatment interaction (F(1, 15) = 0.002, p = 0.963). In line with this, pair-wise post-hoc comparisons revealed no differences between vehicle-treated WT and SOD1^G93A^ mice (WT-VEH = 108.1 ± 32.45, SOD-VEH = 117.6 ± 11.53, p = 0.998), nor between caffeine-treated WT and SOD1^G93A^ animals (WT-CAF = 120.7 ± 40.56, SOD-CAF = 133 ± 17.8, p = 0.998) (Figure 5H).

### Caffeine treatment decreased BDNF levels in the cortex of SOD1^G93A^ mice

Since BDNF is regulated by A_2A_R, the effects of chronic caffeine treatment on cortical and spinal cord BDNF protein levels were assessed in a similar manner to what was done for VEGF.

To assess changes in cortical BDNF protein levels (Figure 6A), a two-way ANOVA of the data obtained by ELISA, revealed a significant main effect of genotype (F(1, 17) = 4.81, p = 0.042), but no significant main effect of treatment (F(1, 17) = 0.178, p = 0.679). Furthermore a significant genotype x treatment interaction (F(1, 17) = 19.2, p < 0.001) was observed. Pair-wise post-hoc comparisons showed that caffeine treatment increased BDNF protein levels in WT mice, when compared with vehicle-treated WT animals (WT vehicle = 20.96 ± 1.403, WT-CAF = 40.24 ± 6.342, p = 0.020). On the other hand, the opposite effect was observed for SOD1^G93A^mice since caffeine treatment led to a significant decrease in BDNF protein levels as compared with vehicle-treated SOD1G93A animals (SOD-VEH = 29.75 ± 4.560, SOD-CAF = 13.85 ± 2.372, p = 0.042). Moreover, a significant difference was observed between caffeine-treated WT and SOD1^G93A^ animals, whereby the latter had significantly lower BDNF protein levels (WT-CAF = 40.24 ± 6.342, SOD-CAF = 13.85 ± 2.372, p = 0.001).

**Figure 6.**
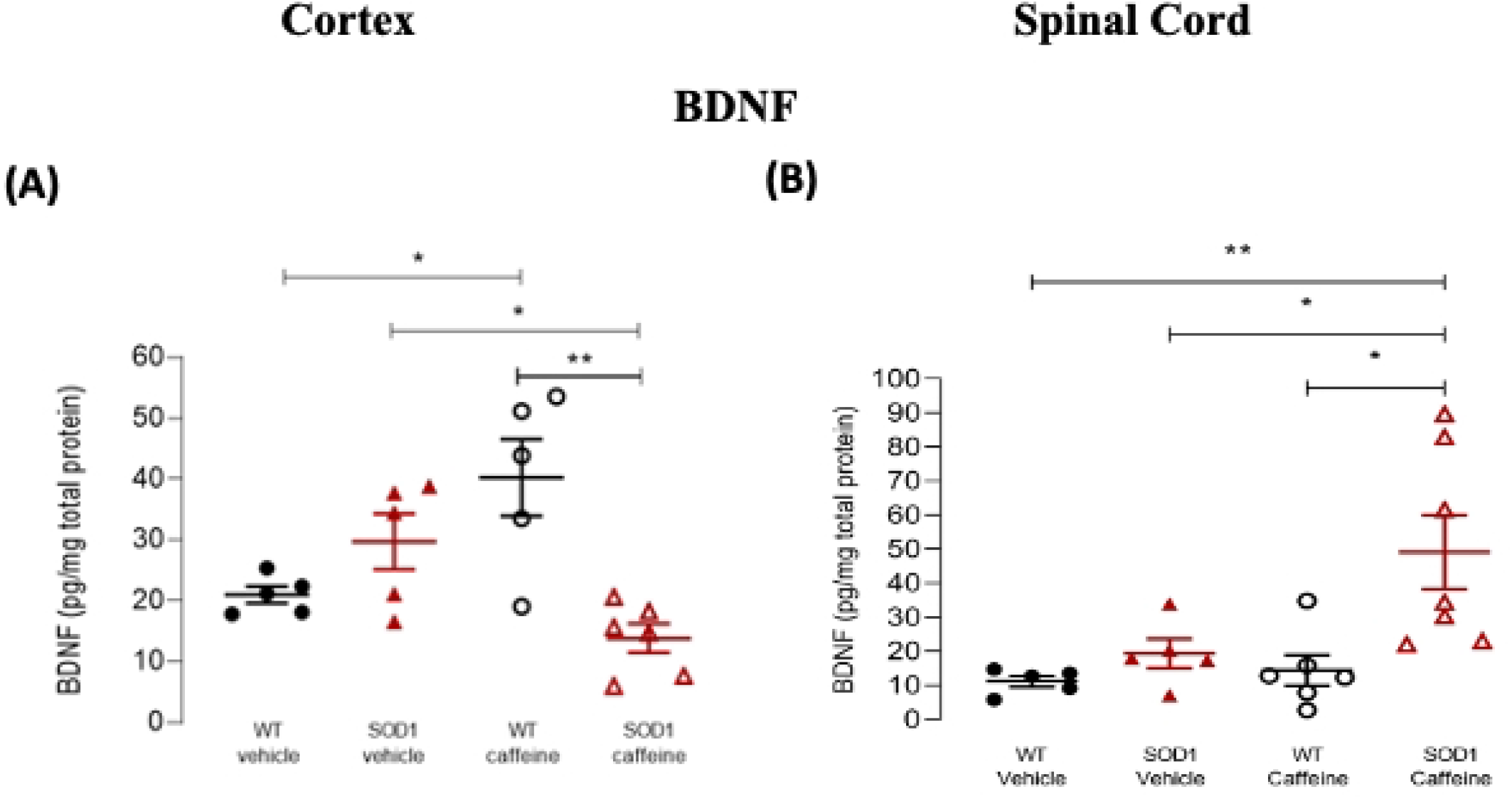
BDNF protein levels in the cortex and spinal cord of WT and SOD1^G93A^ mice treated/non-treated with caffeine. BDNF protein levels in samples from the cortex **(A)** or spinal cord **(B)** of late stage SOD1^G93A^ mice (16-18 weeks old) or age-matched WT mice, treated/non treated with caffeine, as indicated below each data set. The Y-axis represents the BDNF protein levels, determined by ELISA, in pg/mg of total protein. Caffeine administration in the drinking water (see methods) started when animals were 11 weeks old. Data are expressed as mean ± SEM (n = 4-6); *p < 0.05, **p ≤ 0.01, two-way ANOVA with Holm-Sidak correction for multiple comparisons.

Regarding spinal cord BDNF protein levels (Figure 6B), two-way ANOVA found a significant main effect of genotype (F(1, 19) = 8.28, p = 0.010), and a significant main effect of treatment (F(1, 19) = 4.89, p = 0.040), but no significant genotype x treatment interaction (F(1, 19) = 3.18, p = 0.091). Pair-wise post-hoc comparisons revealed that caffeine treatment significantly increased BDNF protein levels in SOD1^G93A^ animals in relation to vehicle-treated SOD1^G93A^ mice (SOD-VEH = 19.34 ± 4.301, SOD-CAF = 49.10 ± 10.82, p = 0.038), and that this increase was also significant when comparing caffeine-treated SOD1^G93A^ mice with caffeine-treated WT mice (WT-CAF = 14.37 ± 4.488, SOD-CAF = 49.10 ± 10.82, p = 0.011). It is important to note, however the high variability in BDNF levels in the spinal cord of SOD1^G93A^ mice treated with caffeine, with 3 out of 6 mice having values close to vehicle-treated mice and the remaining having values clearly above the control levels.

In summary, caffeine treatment lad to a clear decrease in BDNF levels in the cortex of SOD1^G93A^ mice, in clear contrast with what occurs in WT mice. Curiously, this decrease in BDNF levels did not occur in the spinal cord.

### Caffeine treatment modulates A_1_R levels in the cortex of SOD1^G93A^ mice

Lastly, we assessed whether caffeine exposure led to changes in the levels of its receptors, the A_1_R or A_2A_R, and whether differences could be observed between WT and SOD1^G93A^ mice.

Analysis of cortical A_1_R protein levels revealed a significant main effect of genotype (F_(1, 16)_ = 16.8, p < 0.001), and a significant main effect of treatment (F_(1, 16)_ = 16.2, p < 0.001), but no significant genotype x treatment interaction (F_(1, 16)_ = 0.008, p = 0.931). Pair-wise post-hoc comparisons revealed a decrease in A_1_R protein levels vehicle-treated SOD1^G93A^ mice when compared to vehicle-treated WT animals (SOD-VEH = 0.697 ± 0.029, p = 0.045) (Figure7A), corroborating the lower cortical levels of A_1_R previously observed at both pre-symptomatic and symptomatic SOD1^G93A^ mice (Figure 1D). Importantly, this decrease was reversed by chronic caffeine treatment, such that no statistical difference was observed between caffeine-treated SOD1^G93A^ mice and vehicle-treated WT animals (SOD-CAF = 0.994 ± 0.113, p = 0.995). However, prolonged caffeine treatment also caused a significant increase in A_1_R levels in the cortex WT mice (Figure 7A) as previously reported for cortico-hippocampal membranes^37^. When comparing caffeine-treated WT and SOD1^G93A^ mice a significant difference was found, whereby the latter group had lower A_1_R protein levels than the former (WT-CAF = 1.284 ± 0.085, p = 0.045, SOD-CAF = 0.994 ± 0.113, p = 0.045) (Figure 7A).

**Figure 7.**
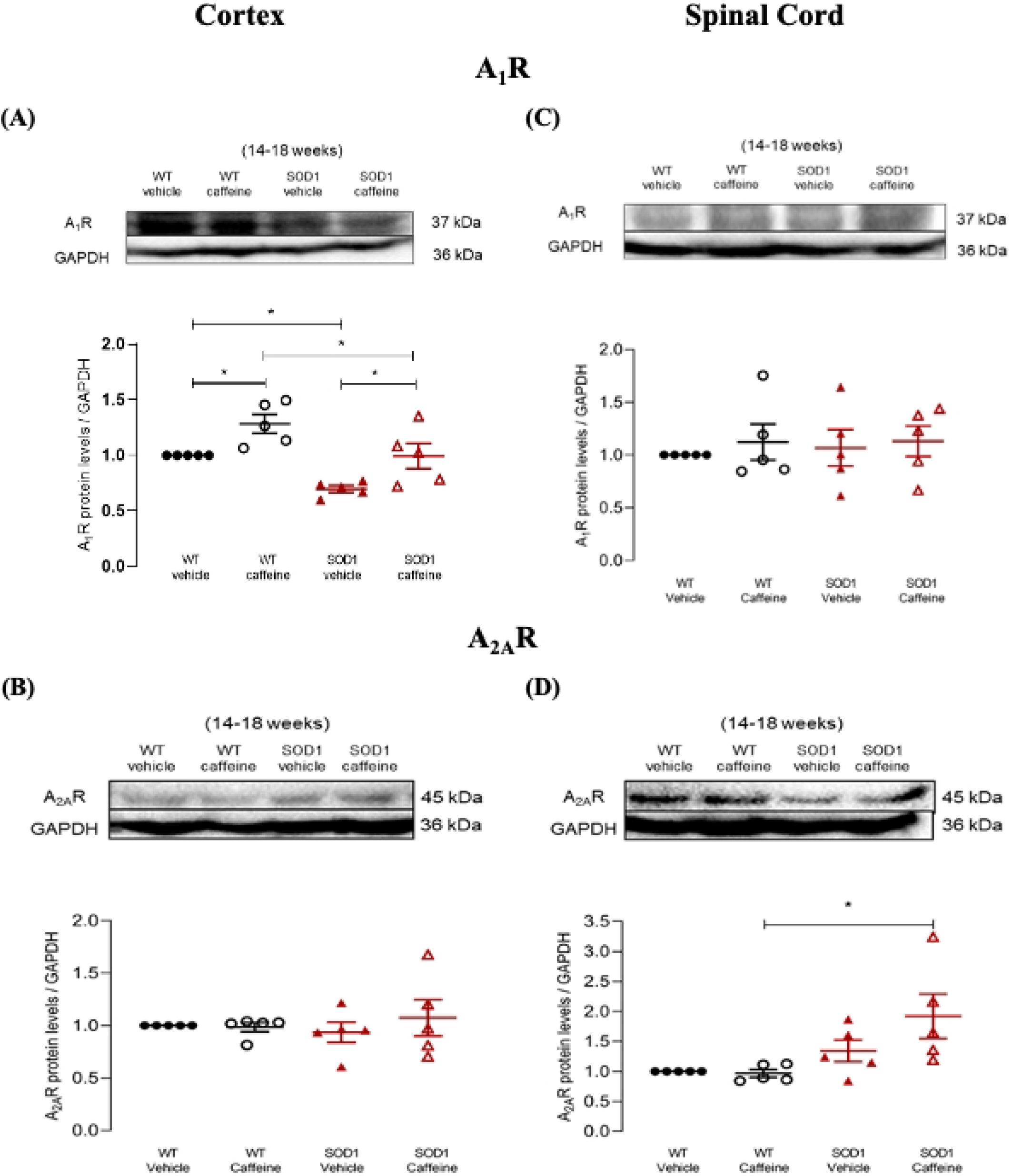
A_1_R and A_2A_R protein levels in the cortex and spinal cord of WT and SOD1^G93A^ mice treated/non-treated with caffeine. A_1_R (A, C) and A_2A_R (B, D) protein levels determined by Western Blot in samples from the cortex (A, B) or spinal cord (C, D) of late-stage SOD1^G93A^ mice (16-18 weeks old) or age-matched WT mice treated/non-treated with caffeine, as indicated below each data set. The Y-axis represents the receptor/GAPDH ratios normalized to values in samples from WT mice not treated with caffeine. Caffeine administration in the drinking water (see methods) started when animals were 11 weeks old. Upper panels show representative immunoblots for each condition. Data are expressed as mean ± SEM (n=5 for all conditions); *p < 0.05, two-way ANOVA with Holm-Sidak correction for multiple comparisons.

Regarding cortical A_2A_R protein levels, analysis found no significant main effects of either genotype (F_(1, 16)_ = 0.013, p = 0.911), or treatment (F_(1, 16)_ = 0.373, p = 0.550), nor a significant genotype x treatment interaction (F_(1, 16)_ = 0.566, p = 0.463). Congruently with this, pair-wise post-hoc comparisons revealed no differences between any of the four groups (WT-CAF = 0.986 ± 0.043, SOD-VEH = 0.936 ± 0.097, SOD-CAF = 1.073 ± 0.172, p = 0.981) (Figure 7B).

### Caffeine treatment modulates A_2A_R levels in the spinal cord of SOD1^G93A^ mice

A_1_R protein levels in the spinal cord of animals treated with caffeine are not significantly affected either in WT or in SOD1^G93A^ mice (Figure 7C). Also, there was no significant differences between WT and late stage SOD1^G93A^ mice (Figure 7C), in agreement with what was found in the spinal cord of early-stage SOD1^G93A^ mice (Figure 2C). Indeed, two-way ANOVA did not detect a significant main effect of genotype (F_(1, 16)_ = 0.075, p = 0.788), or treatment (F_(1, 16)_ = 0.429, p = 0.522), nor a significant genotype x treatment interaction (F_(1, 16)_ = 0.044, p = 0.837). Pair-wise post-hoc comparisons revealed no significant differences between any of the four groups (WT-CAF = 1.122 ± 0.170, SOD-VEH = 1.068 ± 0.173, SOD-CAF = 1.131 ± 0.145, p ≥ 0.995 for all comparisons) (Figure 7C).

Analysis of A_2A_R protein levels in the spinal cord revealed a significant main effect of genotype (F_(1, 16)_ = 9.71, p = 0.007), but no significant main effect of treatment (F_(1, 16)_ = 1.73, p = 0.207), nor a significant genotype x treatment interaction (F_(1, 16)_ = 2.18, p = 0.159). Pair-wise post-hoc comparisons only found a statistically significant difference between caffeine-treated WT and SOD1^G93A^ treated animals, whereby caffeine-treated SOD1^G93A^ mice had significantly higher A_2A_R protein levels (WT-CAF = 0.967 ± 0.063, SOD-CAF = 1.920 ± 0.369, p = 0.030) (Figure 7D).

Summarizing, prolonged caffeine treatment led to a significant increase in A_1_R levels in the cerebral cortex of SOD1^G93A^ mice but of similar magnitude of what it did in WT mice. In contrast, in the spinal cord it led to a selective increase in A_2A_R only in SOD1^G93A^ mice.

## Discussion

The present work shows modifications of adenosine receptor protein levels across disease progression in SOD1^G93A^ mice, with a few modifications in VEGF or VEGF receptors levels. We detected a decrease in A_1_R protein levels in the cortex from the pre-symptomatic stages onwards, but no measurable change in the spinal cord. For A_2A_R protein levels, a transient increase was detected in the spinal cord at the pre-symptomatic stage and in the cortex in the early symptomatic stage. For VEGF signaling we could only detect a transient increase in the levels of VEGFB and VEGFR-2 protein levels in the spinal cord of pre-symptomatic mice, a transient decrease in VEGFR-1 levels in the cortex of early symptomatic mice and a decrease in VEGFA levels in late symptomatic SOD1^G93A^ mice, when compared with age-matched WT mice. BDNF levels, which were assessed only in the late symptomatic stage, were unaltered in non-treated SOD1^G93A^ mice as compared with age-matched WT mice.

While comparing changes in adenosine receptor levels with changes in the corresponding transcripts, it becomes clear that both measurements do not always go in the same direction. A poor correlation between mRNA and protein levels has been described to exist in other situations^38,39,40^ and may result from feedback compensatory mechanisms triggered by the changes in receptor signalling. In addition, besides transcription itself, post-transcriptional, translational, and degradative regulation, can also affect protein levels. Since the receptor and its ligand are the very first partners to trigger a response, in the present work we mostly focused on the levels of the proteins rather than on their transcripts.

The adenosine A_1_R, is a well-known inhibitor of glutamate release in most, if not all, CNS synapses, including those from the cortex^41,42^ and spinal cord^43,44^. Indeed, A_1_R are usually considered as neuroprotective^45,46^.Therefore, the early decrease in A_1_R, now detected in the cortex of the SOD^G93A^ mice, together with the decreased ability of A_1_R to attenuate A_2A_R-mediated responses, reported at the neuromuscular junction SOD^G93A^ mice^31^, may contribute to aggravate neurodegeneration. However, other factors are certainly involved in motor neuron degeneration in ALS since selective blockade of adenosine A_1_R, was shown to significantly attenuate motor disease progression in SOD1^G93A^ mice^47^.

The role of A_2A_R in ALS is controversial since both its selective activation48 or blockade^4^ have been shown to delay disease progression in SOD1^G93A^ mice. Absence of influence of A_2A_R agonists and antagonists upon progressive loss of motor skills or survival of SOD1^G93A^ mice have also been reported^47^. Knowing that A_2A_R signaling is exacerbated at the neuromuscular junction of pre-symptomatic SOD1^G93A^ mice^30^, we addressed the possibility that A_2A_R could be also altered in the spinal cord or in the cortex of SOD1^G93A^ mice. We could indeed detect an early but transient enhancement of A_2A_R levels in the cortex, detected at the pre-symptomatic stage, followed by a later enhancement in the spinal cord, which was detected in the early symptomatic stage. Prolonged caffeine consumption further exacerbated the increase in A_2A_R levels in the spinal cord of SOD1^G93A^ mice. The early increase in A_2A_R in the cortex of pre-symptomatic stage, together with the decrease in A_1_R levels, may further contribute to an exacerbated excitatory drive-in cortical neuron at early disease stages. Cortical hyperexcitability preceding the onset of clinical motor symptoms has been reported in familial and sporadic cases of ALS^49,50^, and has been pointed out as having diagnostic utility^50^. Enhanced excitatory drive in cortical^51,52^ and hippocampal neurons at early disease stages has indeed been reported in SOD^G93A^ mice^53,29^. Early hyperexcitability also occurs at the neuromuscular junction of SOD^G93A^ mice^54^, which may result from decreased A_1_R signaling^31^ and enhanced A_2A_R signaling^30^, which may favor BDNF signaling^55^.

Caffeine had no effect on spinal cord A_1_R levels while inducing the expected^37,56^ upregulation of A_1_R levels in the cortex. This may be due to the fact that A_1_R is the predominant adenosine receptor in the cortex^57,27^. In agreement, there was no effect of caffeine on cortical A_2A_R levels, where the A_2A_R expression is not as predominant. On the other hand, in the spinal cord, where A_2A_R levels were already increased in SOD1^G93A^ mice, caffeine further increased those levels. Interestingly, some studies have shown that a strong expression of A_2A_R in spinal motor neurons, being higher than that of A_1_R^58,4^, might have contributed to a more pronounced effect of caffeine on A_2A_R levels in the spinal cord.

In contrast with what has been shown in other animal models of neurodegenerative diseases^59, 60,61^, caffeine exacerbates ALS symptoms in SOD1^G93A^ mice (Potenza et al., 2013). We thus hypothesized that the negative influence of caffeine upon ALS progression could result from a change in adenosine receptor density levels that in turn would affect neurotrophic factor signaling. Since VEGF is downregulated in cells overexpressing SOD1^G93A^ protein^12^, VEGF mRNA is downregulated in the spinal cord of SOD1^G93A^ mice^12^, and VEGF signaling has neuroprotective roles in motor neurons^62,10^, we first addressed the possibility that caffeine could affect VEGF signaling. However, while comparing the levels of VEGF or VEGF receptors in late stage of the SOD1^G93A^ mice treated/ non-treated with caffeine, no significant alterations that could be attributed to caffeine were detected.

A dual role for BDNF upon ALS progression has been highlighted (Pradhan et al., 2019), since the usual neuroprotective action of BDNF seems to be overcome by a negative influence of BDNF in ALS due to overactivation of truncated TrKB receptors in the spinal cord^63,64^. Interestingly, we detected significantly higher levels BDNF as well as of A_2A_R protein in the spinal cord of SOD1^G93A^ mice under caffeine. This agrees with the fact that A_2A_R promote BDNF synthesis^65,66,67^. Exacerbated excitotoxicity and excitability mediated by A_2A_R and BDNF in the spinal cord may thus contribute to the negative influence of caffeine in SOD1^G93A^ mice survival. However, it should also be noted that caffeine led to decreased BDNF levels in the cortex of SOD1^G93A^. In cortical neurons BDNF is usually regarded as neuroprotective^68,69^. Our results may therefore suggest that the negative influence of caffeine in the ALS mouse model results from a gain of BDNF toxic function in the spinal cord together with a loss of BDNF neuroprotective function in the cortex, which cannot be compensated and may even be exacerbated by the caffeine-induced upregulation of A_1_R.

Along disease progression no significant changes in VEGFA or its main receptor, VEGFR-2 were detected, except for a transient increase in VEGFR-2 levels in the spinal cord of pre-symptomatic mice that turns into a decrease in the symptomatic stage. Interestingly, it was also in the spinal cord and in the symptomatic stage that an increase in A_2A_R levels were detected. Whether enhanced VEGFR-2 levels, which were not accompanied by an increase in its putative ‘decoy’ receptor, the VEGFR-1, contribute to exacerbated spinal cord excitability, thus favoring excitotoxicity, or will result from an early homeostatic neuroprotective mechanism, cannot be answered in the present work. VEGFB levels were also increased in the spinal cord of pre-symptomatic SOD1^G93A^ mice in comparison with age-matched WT mice, and this may favor neuroprotection. The neuroprotective role of VEGFB^9^ may however be transient, failing in the symptomatic stage, since VEGFB levels in the spinal cord of symptomatic SOD1^G93A^ mice were significantly lower than those detected in the pre-symptomatic stage. It is likely that the pre-symptomatic increase in VEGFB protein levels in the spinal cord, represent an early compensatory mechanism, that eventually breaks down as the disease progresses. Interestingly, in motor neurons and spinal cord of post-mortem samples from ALS patients, reduced VEGFA and VEGFR-2 expression was detected^13^. A significant decrease in VEGFA in the cortex, but not in the spinal cord, of late stage SOD^G93A^ mice was also detected in the present work.

In summary, we herein put into evidence an unbalanced adenosine-mediated neuromodulation that occurs in a contrasting way in upper and lower motor neuron synapses. Moreover, prolonged caffeine intake has a dual influence upon the expression of the two high affinity adenosine receptors, that is distinct in the cortex and in the spinal cord but somehow correlates with the relative densities of the two receptors in the two brain areas. Prolonged caffeine intake does not seem to influence VEGF signalling but did, however, affect BDNF levels, also in a contrasting way in the cortex and in the spinal cord. Given the negative influence of BDNF upon survival of spinal cord motor neurons, the present results allow to suggest that the negative influence of caffeine upon disease progression in SOD1^G93A^ mice may result from a gain of BDNF toxic function in the spinal cord together with a loss of BDNF neuroprotective function in cortical neurons. By highlighting this possibility our work makes a step forward in the understanding of the role of adenosine receptor ligands in ALS, namely of caffeine, which clearly contrasts with what is known to occur in other neurodegenerative diseases.

## Author Contributions

N.R. and A.M.S. designed the experiments. N.R. performed the experiments, analysed the data, and wrote the manuscript. C.A.V. helped to perform and analyse qRT-PCR and ELISA experiments and revised the manuscript. M.F.F. participated in the statistical analysis and contributed to the writing and revision of the manuscript. S.H.V. helped with animal dissection and revised the manuscript. J.A.R. and A.M.S. revised the manuscript. All authors approved the final version of the manuscript.

## Acknowledgments

The authors thank Iolanda Moreira and Bruno Novais (iMMRodent facility) for their help in the management and upkeep of the mouse colony.

## Legends

**Supplementary Table 1.**
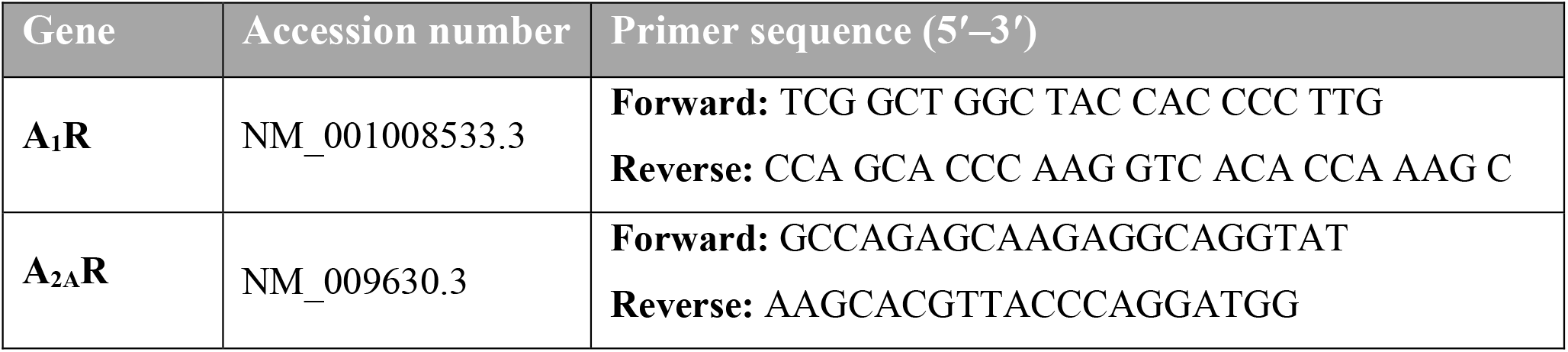
Primers used in the qRT-PCR analysis. The table indicates the gene, the gene accession number and the primer sequence.

